# Inflammatory bowel disease risk gene C1ORF106 regulates actin dynamics in intestinal epithelial cells

**DOI:** 10.1101/2025.03.14.643205

**Authors:** Isabelle Hébert-Milette, Chloé Lévesque, Jean Paquette, Marie-Ève Rivard, Louis Villeneuve, Gabrielle Boucher, Philippe Goyette, Guy Charron, John D. Rioux

## Abstract

**Background and aims:** C1ORF106 has previously been associated with inflammatory bowel diseases (IBD) via large-scale genetic studies. Increased intestinal permeability is a hallmark of IBD and is observed in at-risk individuals prior to the appearance of clinical symptoms. C1ORF106 was previously shown to regulate intestinal barrier permeability through the regulation of adherens junction stability and through the formation of tight junctions, which impacted actin assembly. However, the downstream impact and molecular mechanisms involved in actin regulation by C1ORF106 haven’t been explored. Our study aimed at identifying which pathways involved in intestinal epithelial barrier regulation and F-actin regulation are impacted by C1ORF106 and its IBD-associated variant.

**Methods:** We knocked down (KD) the expression of *C1ORF106* in human colonic epithelial cells and characterized the function of the 333F variant in intestinal epithelial spheroid cultures obtained from patient-derived human induced pluripotent stem cell (hiPSC). We measured barrier permeability and characterized spheroid formation, actin regulation and cell migration though immunofluorescence, western blots and permeability assays.

**Results:** C1ORF106 KD leads to impaired cortical actin belt dynamics and regulation of stress fiber formation, resulting in increased cell constriction, impaired barrier permeability, cell polarity and cell migration. Moreover, we demonstrated that an inhibition of ROCK rescues the actin belt and cell polarity phenotypes in C1ORF106 KD cells, demonstrating that C1ORF106 regulates these phenotypes through a ROCK-dependent mechanism. We also observed an altered nmMYO2-P localization in C1ORF106 KD cells associated with the formation of Vacuolar Apical Compartments (VACs), which are important for 3D epithelial spheroid formation. We observed a similar impact on cell polarity in intestinal epithelial spheroids obtained from hiPSC carrying the 333F variant, providing additional support that this pathway is involved in disease development.

**Conclusion:** We provide insights into the molecular mechanisms by which C1ORF106 controls actin dynamics to regulate intestinal epithelial integrity.

**summary:** C1ORF106 and its inflammatory bowel disease-associated genetic variant regulate intestinal barrier permeability through the regulation of tight junction formation and cell polarity in epithelial cells. This regulation is associated with altered F-actin dynamics that are ROCK-dependent.

## Introduction

Inflammatory bowel diseases (IBD), including Crohn’s disease (CD) and ulcerative colitis (UC), are characterized by chronic inflammation in the gut resulting in intestinal mucosal damage (1, 2). Mucosal integrity is characterized by an intact intestinal barrier function that prevents the activation of the immune system by limiting the passage of gut microbes and of pathogenic antigens (1, 2). The composition and structure of tight junctions (TJ) and adherens junctions (AJ) that form the apical junctional complex (AJC) control the selective passage of molecules between epithelial cells (2). An increased intestinal permeability in unaffected first-degree relatives of CD patients is associated with an increased risk of developing CD (3), suggesting an important role of the epithelial barrier in IBD establishment. Moreover, increasing evidence indicates that achieving mucosal healing can predict long-term remission in IBD (1), highlighting the importance of epithelial homeostasis in IBD.

It was previously demonstrated that *C1ORF106*, a gene associated with IBD, regulates epithelial permeability and that its IBD-associated 333F rare coding variant impairs protein stability (4, 5). Specifically, C1ORF106 KD impacts on intestinal permeability by inhibiting CYTH1 degradation, which activate ARF6 and alters AJ stability (5). Moreover, the association of the TJ protein ZO-1 to cellular membranes is impaired in epithelial monolayers derived from C1ORF106 KO (Knock-out) primary human colonic cells (6), suggesting that C1ORF106 can regulate epithelial permeability by regulating both AJ and TJ. One important regulator of the intestinal epithelial barrier is the actin cytoskeleton (7). Interestingly, some results suggest that C1ORF106 is implicated in the actin cytoskeleton regulation. Indeed, in Caco-2 BBe cells, C1ORF106 increases F-actin assembly at junctions by recruiting CYTH2 to the cell membranes (8). Moreover, C1ORF106 accumulates at junctions in HCT8 cells overexpressing *C1ORF106* when treated with an inhibitor of Rho Associated Coiled-Coil Containing Protein Kinase (ROCK), a regulator of the actin cytoskeleton, and this accumulation enhances epithelial barrier function (9).

To better understand how C1ORF106 affects epithelial integrity, we investigated the impact of knocking down the expression of C1ORF106 on actin cytoskeleton regulation and identified the molecular pathways involved in this process. We report how decreasing C1ORF106 levels in a human intestinal epithelial model affects the regulation of actin contraction and cell polarity establishment. We observe a ROCK-dependent dysregulation of barrier integrity associated with an impaired F-actin regulation. Moreover, we validate the impact of the IBD-associated coding variant *C1ORF106*-*333F* by demonstrating that intestinal epithelial cultures derived from human induced pluripotent stem cells (hiPSC) heterozygous for the 333F allele have an altered cell polarity establishment. These results highlight the multiple effects of C1ORF106 in intestinal homeostasis and further emphasizes the need to develop novel therapeutic strategies that target epithelial functions to complement existing therapies.

## Material and methods

### Caco-2 cell culture

Caco-2 (HTB-37, ATCC) cells were maintained at 70-80% confluence in Complete Medium (EMEM (Cedarlane), 20% fetal bovine serum (FBS, Sigma), 100U/ml penicillin and streptomycin (P/S; Wisent), and 5µg/ml puromycin (Sigma) at 37°C with 5% CO_2_. Cells were plated on glass coverslips (Life Technologies; coated with 15µg/ml laminin (#cat: 23017015, Gibco) for 15 minutes at room temperature (RT)) or on polytetrafluoroethylene (PET) 12-wells Transwell filters (Corning; coated with 65.3µg/ml laminin (37°C, 5% CO_2_) for 24h) and were differentiated for 21 days in polarized epithelial monolayers. Medium was changed every 2-3 days.

### Lentiviral infections

Cells were infected as described previously in (5). Briefly, C1ORF106 and control shRNA vectors (TRCN0000140233 and SHC001V, Sigma, MISSION) were added in a ratio 2:2:1 to the lentiviral packaging and envelope vectors (Sigma, MISSION) and transfected into HEK293T cells by calcium phosphate precipitation (Open Biosystems protocol). After 48h, lentivirus-containing medium was collected and filtered through a 0.45µm filter and lentiviral particles were titrated with the QuickTiter Lentivirus-Associated p24 Titer Kit (Cell Biolabs). Caco-2 cells were plated at 50% confluence 24h before the infection with a lentiviral particle MOI of ∼10 in minimal medium (EMEM, 1% FBS, 8µg/ml polybrene). Medium was changed after 24h and cells were selected with puromycin (10µg/ml) after 48h. C1ORF106 KD level was confirmed by western blot and qPCR.

### Cell number quantification

Cells were differentiated in 6-well plates (ThermoFisher), washed and treated with 2mM EGTA pH 7.4 for 30 minutes (37°C, 5% CO_2_). Viable cells were counted using Trypan Blue and a hemocytometer. MTT assay: cells were differentiated in 12-well plates (VWR), washed and treated with 0.225mg/ml 3-(4,5-Dimethylthiazol-2-yl)-2,5-Diphenyltetrazolium Bromide (MTT) for 1h30 (37°C, 5% CO_2_). MTT was removed, and the reaction stopped with DMSO. Absorbance was measured at 570 and 650nm (reference wavelength) with the Biotek spectrometer and reported to a standard curve. AlamarBlue assay: differentiated cells were incubated for 3h30 (37°C, 5% CO_2_) with the AlamarBlue™ Cell Viability Reagent (ThermoFisher). Fluorescence was measured with an excitation/emission (570nm/600nm) wavelength with the Infinite 1000 Pro (Tecan) and was reported to a standard curve.

### Confocal Immunofluorescence

Cells were differentiated on laminin-coated glass coverslips, washed with PBS pH 7.2, fixed for 20 minutes with 2% paraformaldehyde, washed, and permeabilized (0.5% Triton X-100, 2% donkey serum in PBS pH 7.2) for 30 minutes at RT. Cells were incubated with primary antibodies (Table S1) in antibody buffer (0.05% Triton X-100, 0.5% donkey serum in PBS pH 7.2) overnight at 4°C, washed, incubated with secondary antibodies for 4h at RT and washed. The coverslips were mounted using 0.4% DABCO (Sigma) in glycerol. Images were acquired as Z-stacks with a LSM 710 confocal microscope using a 63x/1.4 oil Plan-Apochromat objective (Carl Zeiss), and were processed using the ZEN 2012 software (Carl Zeiss, Blue edition). Focal adhesions were quantified in images taken with the INcell Analyzer 6000 (GE Healthcare Life Sciences) and analyzed with the INcell Developer. The images were segmented and the adhesions of less than 0.099µm^2^ were excluded (nonspecific signal).

### TEER measurement

Cells were differentiated on Transwell filters. TEER was measured with the Epithelial ohm meter EVOM2 (World Precision Instruments). The resistance value (ohms (Ω)/cm^2^) was obtained by subtracting the blank and multiplying by the growth surface area (1.12cm^2^) of the filter.

### Calcium switch

Cells were differentiated on laminin-coated Transwells or on laminin-coated coverslips, treated with 5mM EGTA for 3h (37°C, 5% CO_2_), washed and incubated in complete medium.

### F/G-actin ratio

Cells were differentiated on laminin-coated coverslips. Cells were washed, lysed in the Actin stabilization solution (0.1M PIPES pH 6.9, 30% glycerol, 1% Triton X-100, 1mM EGTA, 1mM MgSO_4_, 1mM ATP, 5% DMSO, protease and phosphatase inhibitors), incubated on ice for 10 minutes and centrifuged at 16 000 x g, 4°C for 75 minutes. The supernatants (G-actin) were collected. The pellets were resuspended in Actin depolarization solution (0.1M PIPES pH 6.9, 1mM MgSO_4_, 10mM CaCl_2_, 5µM cytochalasin D, protease and phosphatase inhibitors), incubated on ice for 1h and centrifuged at 16 000 x g, at 4°C for 75 minutes. The supernatants (F-actin) were collected. Actin fractions were quantified by immunoblotting using a β-actin antibody.

### Immunoblotting

Proteins were extracted in lysis buffer (50mM Tris-HCl pH 7.6; 150mM NaCl, 1mM EDTA, 1% NP-40, 1% Triton X-100, protease and phosphatase inhibitors), and were centrifuged at 16 000 x g at 4°C for 15 minutes. The supernatant protein concentrations were determined using the Pierce BCA protein assay (ThermoFisher). Proteins were boiled at 95°C for 10 minutes in Laemmli buffer (Bio-Rad), separated by electrophoresis on a denaturing polyacrylamide gel and transferred on a nitrocellulose membrane (Bio-Rad) for 2h. Membranes were blocked overnight at 4°C in Tris-buffered saline with 0.1% Tween (TBST) with 5% low-fat milk, incubated for 2h at RT with primary antibodies (Table S1), washed in TBST and incubated with HRP-conjugated antibodies (Abcam, 0.5µg/ml) for 2h at RT. Membranes were washed and incubated with the Western Blot Lightning Plus-ECL reagents (Perkin Elmer). Vinculin was used as loading control. Band intensity was quantified with ImageJ.

### Caco-2 spheroid culture and treatments

45µl of Matrigel (#cat: 354234, Corning) was polymerized for 10 minutes at 37°C in cell imaging coverglass (Eppendorf) before seeding 10 000 cells in complete medium with 2% matrigel. Spheroids were grown for 7 days (37°C, 5% CO_2_) and counted using the cell counter plugin of ImageJ. Confocal immunofluorescence was performed with a permeabilization step of 1h. To assess spheroid permeability, spheroids were incubated with a 0.3kDa fluorescent molecule (2µg/ml, FITC, Sigma-Aldrich) for 30 minutes and imaged with a LSM 710 confocal microscope (Zeiss) using an 10x/0.3 Plan Neo-Fluar objective in an environmental chamber (37°C, 5% CO_2_). The fluorescence intensity was measured with ImageJ and the ratio of [inside spheroids-background (cell fluorescence)]/[outside spheroids-background] was calculated. Spheroids were considered permeable if the fluorescence in the lumen (for spheroids with a lumen) or in cellular junctions (for spheroids without a lumen) was higher than the intracellular fluorescence (background). In the forskolin experiments, spheroids were treated with forskolin (10µM, StemCell Technologies) for 5h. The same spheroids were imaged before and after the treatment. Spheroid area was assessed with ImageJ.

### Migration assay

Cells were differentiated in two-well silicone inserts (#cat: 80209, Ibidi) placed on laminin-coated coverglass. At day 20, serum was removed. At day 21, inserts were removed, cells were washed and fixed for immunofluorescence. For live fluorescence, cells were washed in Phenol red free medium (PRFM) (DMEM without phenol red, 100U/ml P/S and 5µg/ml puromycin), stained for F-actin (Cellmask^TM^) for 1h in PRFM, washed, stained for nucleus (Hoechst) for 20 minutes in PRFM (Table S1) and washed before inserts were removed. Pictures were taken every 20 minutes with a LSM 710 confocal microscope (Zeiss) using an 10x/0.3 Plan Neo-Fluar objective in an environmental chamber (37°C, 5% CO_2_). 10 cells per well (distributed evenly across both sides of the migration front) were analysed with ImageJ Manual Tracking plugin. The distance travelled was determined by subtracting the cell positions at a given time to their initial positions. The directionality was defined as path length (hypotenuse of the distance traveled)/total path length (sum of hypotenuses between each time points).

### ROCK and nmMYO2 inhibition

nmMYO2 inhibition: differentiated cells were challenged with blebbistatin (100µM, Sigma) for 12h and fixed for confocal immunofluorescence. ROCK inhibition: Y-27632 (Abcam, 40µM) was added to the complete medium after the EGTA treatment. During the ROCK inhibition experiment in spheroids, spheroids were grown with Y-27632 (20µM) in the medium.

### hiPSC reprogrammation, characterization, differentiation and spheroid culture

hiPSC were reprogrammed and differentiated as described in (10). Briefly, using whole exome sequencing (WES) data (11) we selected four lymphoblastoid cell lines (LCL) from the NIDDK IBDGC Repository based on their genotype status for the C1ORF106-333^Y/Y^ (rs41313912): two of them (females diagnosed with CD) were homozygous carriers of the wild-type allele (C1ORF106-333^Y/Y^) allele and two (one female and one male diagnosed with UC) were carriers of C1ORF106-333F allele (C1ORF106-333^Y/F^). In addition, these lines were evaluated in the WES data for the presence of other IBD-associated non-synonymous coding variant in known causal genes (NOD2, IL23R, ATG16L1, RNF186, IFIH1, GPR65, CARD9 and MST1). Control-1 was found to be a heterozygote carrier for ATG16L1-T300A and GPR65-I231L, Control-2 did no carry any of the tested variants, C1ORF106-333^Y/F^-1 was heterozygote for ATG16L1-T300A and MST1-R703C, and C1ORF106-333^Y/F^-2 was homozygote for for ATG16L1. The LCL were reprogrammed in hiPSC by nucleofection with four episomal reprogramming plasmids (pCE-hUL, pCE-hSK, pCE-hOCT3/4, pCE-mp53DD; a kind gift from Shinya Yamanaka (Addgene plasmids #41813, 41814, 41855, 41856)(12)). TRA-1-81 positive colonies were selected. The expression of pluripotency markers (NANOG, SOX2 and POU5F1) and the absence of genomic integration of the reprogramming plasmids were confirmed by RT-qPCR. Genotype status for the C1ORF106--333^Y/F^ (rs41313912) variant was confirmed by Sanger sequencing (Génome Québec). Chromosomes were analyzed by Karyostat Assay (Thermofisher). The capacity to form the three germ layers was assessed using the Human Pluripotent Stem Cell Functional Identification Kit (R&D Systems). Polarized intestinal epithelial monolayers were generated through a conversion to definitive endoderm (DE) using the STEMdiff Definitive Endoderm kit (StemCell Technologies) and the formation of hindgut (HG) structures induced by 3µM CHIR99021 and 250ng/ml FGF4 for 3-4 days. HG structures were manually harvested, centrifuged for 5 minutes at 100 x g, embedded in Geltrex (ThermoFisher, #cat: A1413202) and cultured for 15 days in Intestinal Growth Medium (IGM; Advanced DMEM/F12, Glutamax, P/S, N2, B27 (ThermoFisher), human recombinant inducers rhEGF (100ng/ml, Cedarlane), rhNoggin (100ng/ml, Millipore-Sigma) and rhR-Spondin-1 (500ng/ml, Millipore-Sigma)), with medium change every 2-3 days. Spheroids were cultivated in IGM and dissociated every 6-8 days to mature into 3D intestinal epithelial spheroids. Spheroids were recovered in basal media, spotted on the surface of the Geltrex-coated chambered glass coverslips (Ibidi) and were cultivated for 10 days in IGM with 4% Geltrex for characterization.

### Statistical analysis

Unless specified otherwise, each experiment was realized in at least three biological replicates. Statistical analyses were performed on the log scale to account for log-normal distribution and multiplicative effects. The technical replicates were pooled before the analyses. Unpaired t-test with Welch’s correction, paired t-test or one sample t-test were used as described in the figure legends. The mean and standard error (SEM) were calculated and transformed back as geometric mean to the original scale.

### Ethics statement

Experiments with human cell lines were approved by the ethics committee of the Montreal Heart Institute, known as the Comité d’éthique de la recherche et du développement des nouvelles technologies (CERDNT; protocol number 2005-23; 05-813). Written informed consent was obtained for all subjects.

## Results

### C1ORF106 regulates the actin cytoskeleton

As C1ORF106 has been shown to regulate both AJ and TJ (5, 6), we wanted to elucidate the potential mechanistic links between these to better understand its role in susceptibility to IBD. To do so, we knocked down (KD) the expression of the endogenous C1ORF106 protein by ∼73% in Caco-2 cells (herein named C1ORF106 KD cells) (Fig. S1A) as previously reported(5). After 21 days of differentiation, C1ORF106 KD cells appeared smaller than control cells (Fig. 1B). In fact, we found that there were twice as many viable cells by surface area in C1ORF106 KD cultures compared to controls as measured by cell count (1.99-fold change, *P=*.059; Fig. 1A) and viable cells assessment using MTT (1.88-fold change, *P=*.027; Fig. S1B) and AlamarBlue assays (1.82-fold change, *P=*.031; Fig. S1C). Confocal immunofluorescence (Fig. S1D) revealed that there was an equivalent reduction in diameter at the apical and basal sides, with an apparent increase in cell height in C1ORF106 KD cells but that there was no evidence of altered contact inhibition or cell extrusions, suggesting that reduced levels of C1ORF106 in intestinal epithelial cells (IEC) can lead to an altered cell shape.

**Figure 1:**
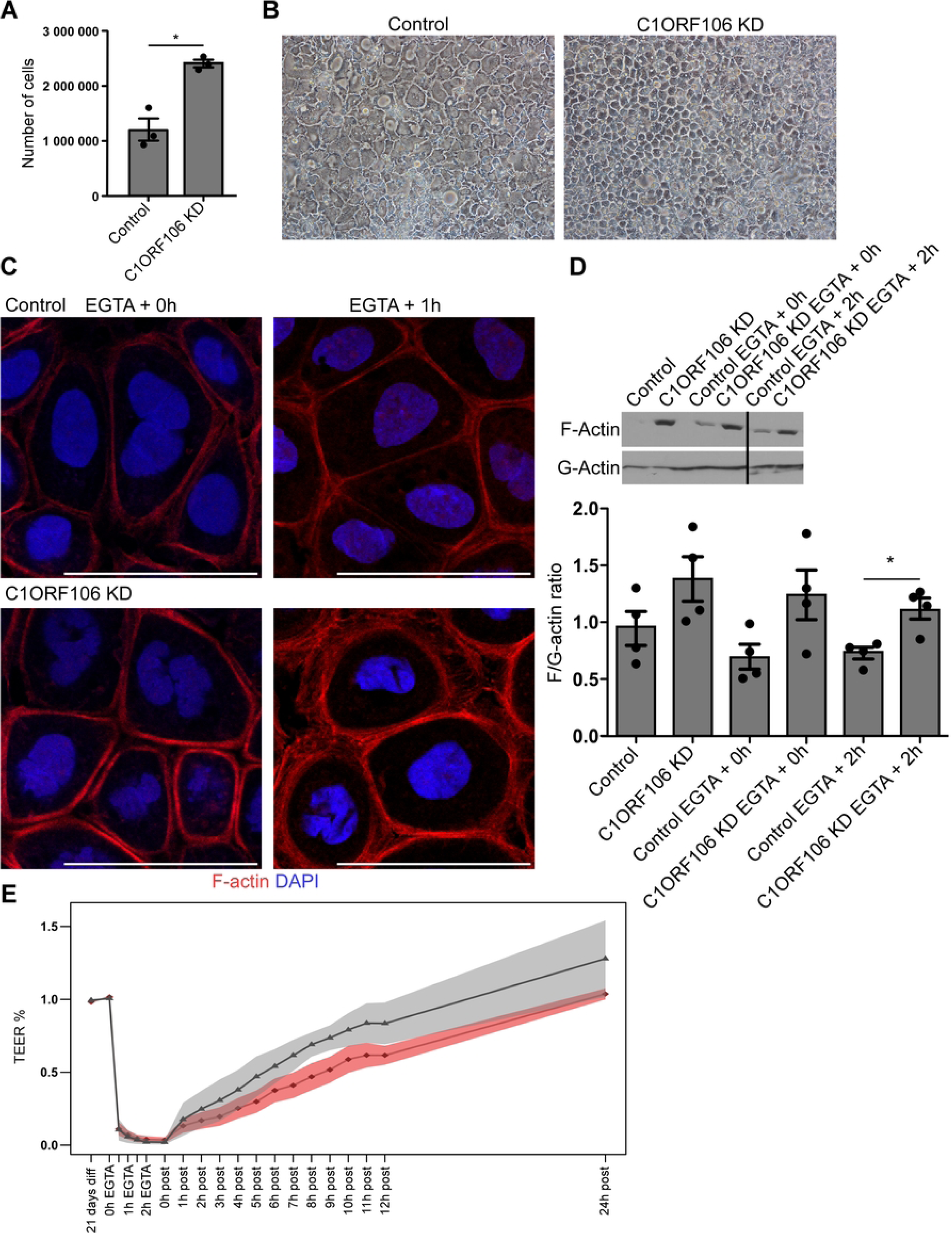
C1ORF106 regulates actin belt contraction. C1ORF106 KD and control Caco-2 cells were differentiated (A-E); treated with EGTA for 3h and allowed to recover for the indicated time (C-E). A) Number of viable cells counted with an hemacytometer. Geometric mean ± SEM (n=3). Unpaired t-test with Welch’s correction. \**P<.*05. B) Brightfield images. C) Confocal immunofluorescence of F-actin (red) and DAPI (blue). Scale bar=50µm. D-E) Geometric mean + SEM. Results are from 2(D) or 3(E) independent experiments. D) F/G-actin ratio. Paired t-test. \**P<*.05., n=4. E) Percentage of TEER recovery. Grey=Control. Red=C1ORF106 KD. Welch two sample t-test. *P<*.05.

Increased actin contractility can pull on cell membranes, which drives cell constriction and impacts on cell shape (13, 14). A dysregulation in the actin belt contraction can alter AJ and TJ regulation by pulling on junctions (14, 15). Notably, we observed in multiple experiments that the F-actin staining appeared stronger in C1ORF106 KD cells than in controls, suggesting an increased actin belt contraction (Fig. 1C). To investigate whether lower levels of C1ORF106 would affect the actin belt during junction formation, we treated our cells with EGTA for 3h to induce AJC internalization and enabled the junctions to reform by placing cells in complete medium. During this dynamic process, actin belt contraction is important for junction destabilization and re-formation (16, 17). Strikingly, the actin belt was thicker in C1ORF106 KD than in controls 1h of recovery after the EGTA treatment (Fig. 1C). As actin filaments (F-actin) are linear polymers of globular actin (G-actin) subunits, we separated these two actin fractions by centrifugation and quantified their abundance. We observed a 1.62-fold (*P<*.05) increase in F/G-actin ratio in C1ORF106 KD cells compared to controls 2h after the EGTA treatment (Fig. 1D), supporting an increased amount of actin structures in these cells. Of note, although not significant (*P=*.089), the trend was also observed at an earlier timepoint (EGTA + 0h), suggesting that the actin dysregulation is present earlier in the experiment. To determine the impact of these changes on barrier permeability, we measured transepithelial electrical resistance (TEER) after the EGTA treatment, and observed that the recovery was slower in C1ORF106 KD cells (5.14%/h) than in controls (7.68%/h) (*P=*.043) (Fig. 1E and S2A), concordant with previous results(5). Moreover, while ZO-1 and F-actin appeared to be colocalized in controls during recovery post-EGTA treatment, this was less evident in C1ORF106 KD cells (Fig. S2B), suggesting that TJ formation is impaired in these cells. These results suggest an impaired epithelial barrier recovery in C1ORF106 KD cells that could be associated with an altered actin regulation.

On the basal side, cell constriction is modulated by the contraction of stress fibers, which are actin bundles attached to focal adhesions (13, 18). Non-muscle myosin-2 (nmMYO2) reorients stress fibers to regulate actin bundle thickness (18). We assessed the localization of F-actin and paxillin, a focal adhesion protein, and observed that the F-actin fluorescence is higher on the basal side of C1ORF106 KD cells, suggesting a dysregulated stress fiber reorganization (Fig. S3A). This is consistent with our observation of a 2.24-fold greater F/G-actin ratio at steady state in C1ORF106 KD cells compared to controls (*P=*.062) (Fig. 1D, S3B). However, no significant alteration was observed in the number or size of focal adhesion sites in C1ORF106 KD cells (Fig. S3C-D), suggesting that the stress fiber phenotype is not associated with focal adhesion regulation. Stress fibers are important for multiple cellular processes including the transmission of the outside-in polarity cue coming from the extracellular matrix (19, 20). An impaired stress fiber regulation could potentially impair signal transmission and cell polarity.

### Cell polarity establishment is impaired in C1ORF106 KD cells

An increased actin contractility in epithelial cells impairs cell polarity establishment, which controls the plasma membrane protein localization (21). We grew our cells into 3D spheroids to evaluate the impact of C1ORF106 KD on cell polarity. We assessed cell polarity in a 3D model as this recapitulates the steps of organ morphogenesis and is therefore more representative of cell polarity establishment than 2D models (22). While 74.6% of spheroids formed by control cells had a defined lumen, this number was lower (35.1%) in C1ORF106 KD cells (Fig. 2A-B) (*P<*.05). As expected, the apical protein marker ezrin was observed on the luminal side of these spheroids, indicating appropriate cell polarity (Fig. 2C). The remaining spheroids had either an inverted polarity (0.1% in controls; 3.4% in C1ORF106 KD) with ezrin located on the external side of spheroids or lacked a lumen (25.3% in controls; 61.4% in C1ORF106 KD) (Fig. 2A-C).

**Figure 2:**
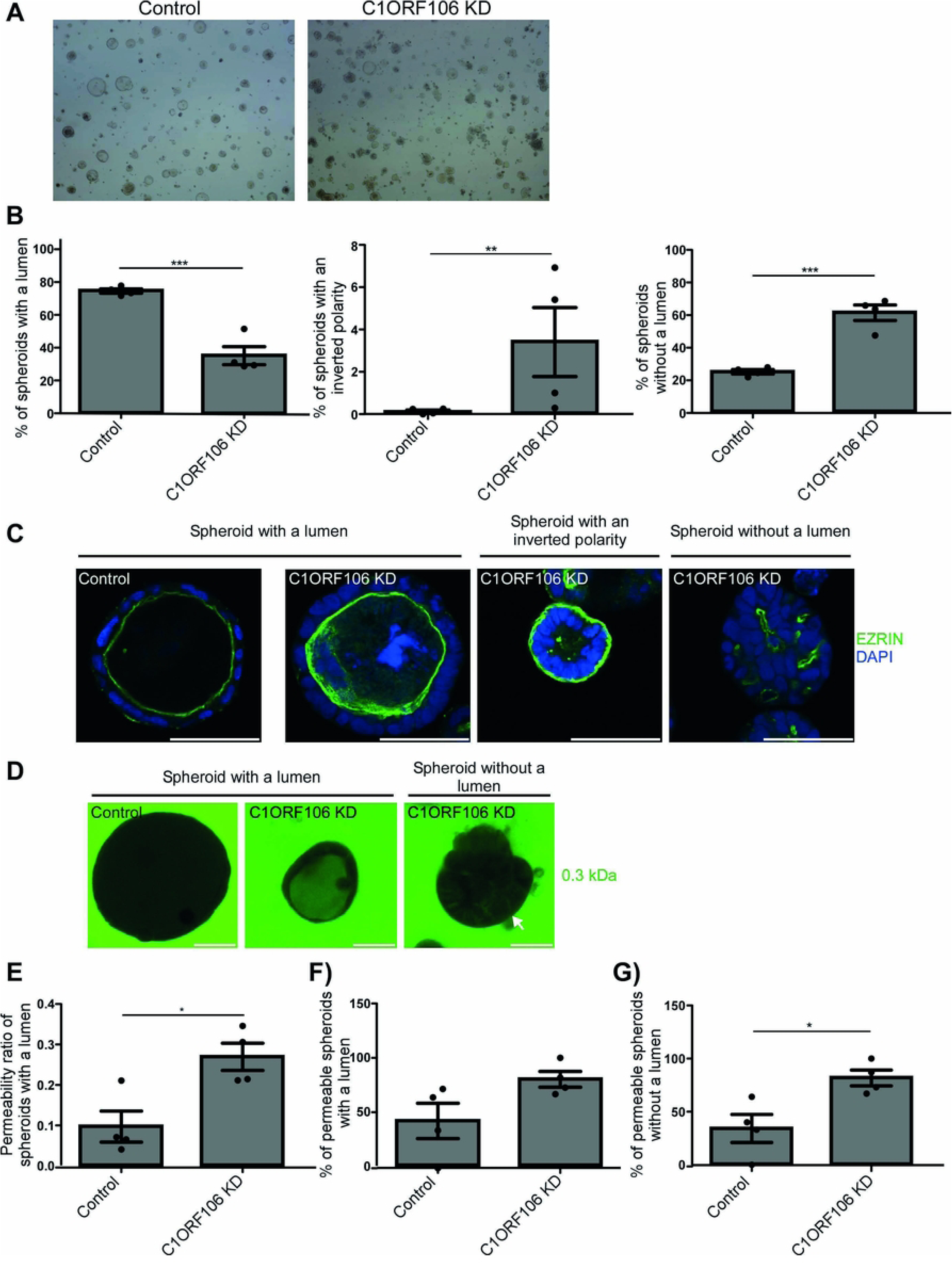
C1ORF106 KD impairs spheroid formation. C1ORF106 KD and control Caco-2 cells were grown into spheroids. Geometric mean ± SEM. A, B) Brightfield images and quantification of the proportion of each type of spheroids. n=4. \**P<*.05; \*\**P<*.005. Paired t-test. C) Confocal immunofluorescence of Ezrin (green) and DAPI (blue). Scale bar=50µm. D-G) Spheroids were incubated for 30 minutes with 2μg/ml FITC (0.3kDa fluorescent molecule). Images (live microscopy) and quantification of spheroid permeability. D) FITC (Green). Scale bar=50µm. E) The fluorescence intensity in spheroids with a lumen was measured as described in the material and method section. Paired t-test. **\****P<*.05. F-G) Proportion of permeable spheroids with a lumen (F) and without a lumen (G) (*P<*.071). Unpaired t-test with Welch’s correction.

In addition to impairing appropriate spheroid polarization, we observed that C1ORF106 KD also increased spheroid permeability. We incubated the spheroids with small fluorescent molecules for 30 minutes and observed that fluorescence intensity is 2.77-fold higher (*P<*.05) inside C1ORF106 KD spheroids with a lumen than inside controls (Fig. 2D-E). Moreover, the proportion of permeable spheroids was greater in C1ORF106 KD than in controls (spheroids with a lumen: 1.91-fold; spheroids without a lumen: 2.38-fold (*P=*.071)) (Fig. 2F-G), suggesting an impaired epithelial barrier integrity.

To better characterize these different spheroid types, we treated them with forskolin, an activator of CFTR-mediated anion and fluid transport, which is unidirectional in polarized cells and can be used to show the polarity of the spheroids (23). We quantified the change in surface area of the spheroids, which is an approximation of the impact on the spheroid volume. Strikingly, by following the same spheroids throughout the experiment, we observed that some of the spheroids without a lumen formed one during the forskolin treatment (Table S2). The proportion of spheroids without a lumen decreased (14.19% in controls; 16.18% in C1ORF106 KD cells; Fig. 3A-B), while the proportion of spheroids with a lumen increased (14.42% in controls; 19.54% in C1ORF106 KD cells; Fig. 3A,C). The forskolin treatment also reduced by 1.64% the percentage of spheroids with an inverted polarity in C1ORF106 KD cells, while these spheroids remained very rare in controls (Fig. 3D). These results suggest that C1orf106 KD cells have the potential for forming well developed spheroids albeit with less success than control cells. Interestingly, we observed that the spheroids reacted differently to forskolin depending on the spheroid type (Videos S1-S3). The ratio of area increase in control and C1ORF106 KD spheroids that started the experiment with a lumen (1.334 and 1.428-fold, respectively) was similar to the area increase in spheroids that started the experiment without a lumen, but formed one (1.402- and 1.438-fold, respectively) (Fig. 3E). However, these two groups of spheroids were significantly different from the spheroids that remained without a lumen after the treatment, which had an area increase of only 1.129-fold for controls and 1.096-fold for C1ORF106 KD (Fig. 3E). These results suggest that the spheroids must first form a lumen before activating CFTR-mediated transport and that C1ORF106 KD spheroids are behaving similarly than controls at the same stage of spheroid formation. The forskolin treatment also reduced spheroid permeability (Fig. 3F-H), suggesting that it promotes epithelial barrier development. Taken together, these results suggest that there are several steps in spheroid formation (Fig. 3I), that the forskolin treatment promotes spheroid progression through these steps and that C1ORF106 KD spheroid formation seems delayed compared to controls.

**Figure 3:**
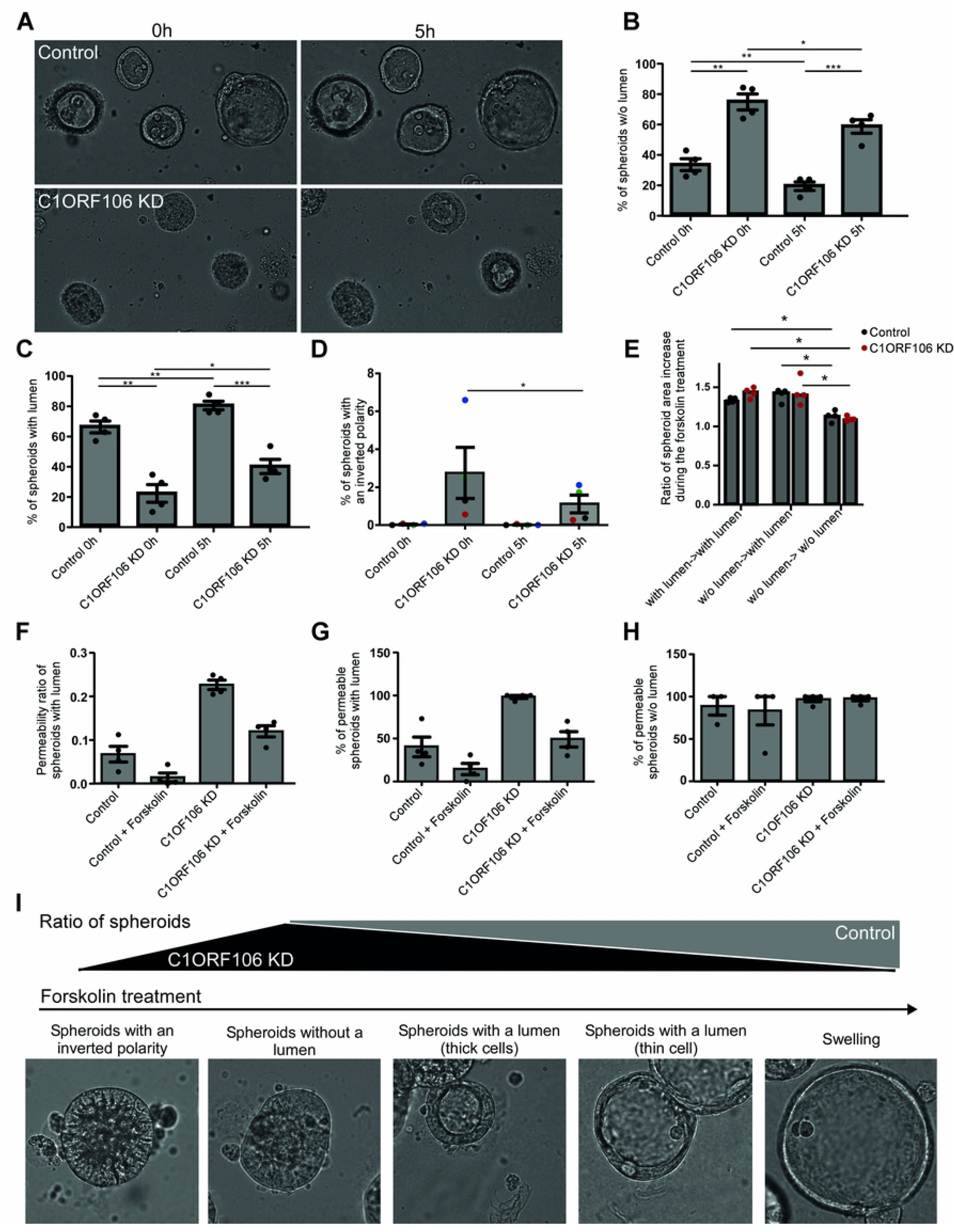
Progression of spheroid formation in C1ORF106 KD cells. C1ORF106 KD and control Caco-2 cells were grown into spheroids and treated with forskolin for 5h. Geometric mean ± SEM. A-D) Brightfield images and quantification of each type of spheroid before and after the treatment. Paired t-test (treated vs non treated) and Unpaired t-test with Welch’s correction (Control vs C1ORF106 KD) \**P<*.05; \*\**P<*.01; \*\*\**P<*.001. Spheroids without a lumen (B); with a lumen (C) and with an inverted polarity (D). D) Dot colors represent each biological replicate. E) Spheroid area increase during the treatment. For each well, spheroids were grouped depending on the spheroid type before and after the experiment. Ten spheroids per group were analyzed. Paired t-test. \*\**P<*.01; \*\*\**P<*.005. F-H) Quantification of spheroid permeability. Paired t-test. n=4. F) Ratio of fluorescence intensity in spheroids with a lumen (C1ORF106 KD treated vs not treated: *P=*.023). G-H) Percentage of permeable spheroids G) with a lumen (C1ORF106 KD treated vs not treated: *P=*.029) and H) without a lumen. I) Observed steps of spheroid formation.

One hypothesis of spheroid formation is that it happens via the delivery of vacuolar apical compartments (VACs) from the outside to the inside of the spheroids (24). VACs are coated with F-actin and are responsible for the transport of apical (e.g. ezrin) and junctional proteins to the apical side (25, 26). Interestingly, we observed in C1ORF106 KD cells an accumulation of F-actin-coated vesicles colocalized with nmMYO2 phosphorylated on the S19 of the myosin light chain, a staining specific to VACs (26), on the basal side of the cells. This is not the typical location for F-actin and nmMYO2 structures, as demonstrated by the absence of nmMYO2-P signal in the control (Fig. 4A). Importantly, the phosphorylation of nmMYO2 is essential for the maturation of VACs and RAB11, which regulates multiple recycling pathways, is recruited downstream of nmMYO2 action for the final maturation step in RAB11-tubular recycling vesicles (24, 26, 27). Strikingly, even though controls showed the presence of round- and tubular-RAB11 vesicles, C1ORF106 KD cells showed almost exclusively round-RAB11 vesicles (Fig. 4B-C), suggesting that the vesicle maturation defect is specific to the tubular-RAB11 recycling pathway. RAB11 didn’t colocalize with VACs in C1ORF106 KD cells (Fig. 4B), suggesting that the maturation of VACs is impaired before their recruitment, with potential impacts on protein delivery to the apical side and cell polarity establishment. Moreover, we observed that RAB11 vesicles seemed more evenly distributed within cells in C1ORF106 KD as compared to control cells (Fig. 4C), suggesting that the localization of RAB11 vesicles might be impacted by C1ORF106, further supporting an alteration in vesicular trafficking and protein delivery in C1ORF106 KD cells.

**Figure 4:**
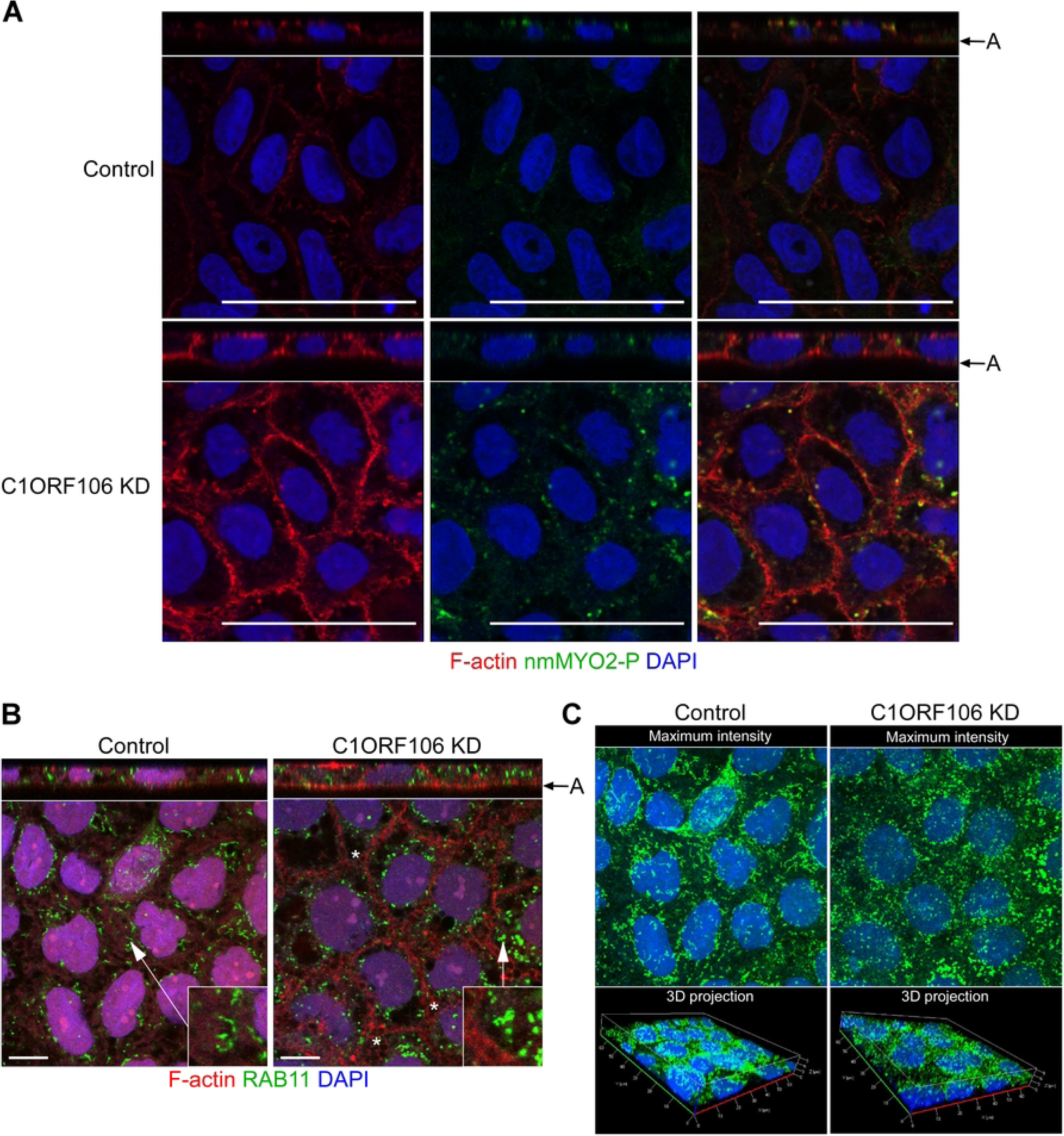
C1ORF106 KD impairs VACs formation. Confocal immunofluorescence of the basal side of differentiated C1ORF106 KD and control Caco-2 cells with DAPI (blue), F-actin (red) and A) nmMYO2-P (Green) (Scale bar=50µm), and B) RAB11 (green) (Scale bar=10µm. Zoom=250%; * = examples of VACs. A=Apical side). C) Maximum intensity projection (top) and 3D projection (bottom) of RAB11 (green) and DAPI (blue).

### C1ORF106 regulates cell migration

As actin dynamics and cell polarity regulate cell migration, we studied how C1ORF106 KD affects cell migration by differentiating our cells with migration inserts for 21 days and arresting cell proliferation by removing serum from the medium for 24h. Inserts were then removed to simulate a wound and cell migration was followed via live confocal microscopy. Following measurements of the distance travelled by cells, we observed that most of the migration happened in the first 4 hours of the experiment for both controls and C1ORF106 KD cells. We observed a higher migration rate in both the x-axis (1.66-fold) and the y-axis in C1ORF106 KD cells compared to control cells (Fig. S4A-B). This increased migration didn’t affect the number of times cells changed direction (Fig. S4C), indicating that they can migrate persistently, but that they do not migrate coordinately in the axis of the migration front. Finally, we noticed that the structure of the migration front in C1ORF106 KD cells is altered as the actin front seems diffused (Fig. S4D). These results suggest that epithelial reorganization regulation following injury is impaired in C1ORF106 KD cells, which impacts on cell migration and tissue repair.

### The C1ORF106 impact on actin cytoskeleton is ROCK dependent

We then determined which actin-regulating molecular pathways are affected by C1ORF106. The actin belt and stress fibers are regulated by multiple pathways including Ras Homolog Family Member A (RhoA) and its effector ROCK, which regulate the phosphorylation and activation of nmMYO2 and stimulate the contraction of actin structures (15, 19). We hypothesized that inhibiting ROCK would correct some of the defects observed in C1ORF106 KD cells. We added a ROCK inhibitor (Y-27632) during the calcium switch recovery. As expected given the role of ROCK and nmMYO2 in junction formation (7), this decreased the actin staining in controls. In comparison, this decreased the actin belt thickness in C1ORF106 KD cells to a level similar to the controls, effectively rescuing the phenotype (Fig. 5A). Moreover, the F/G-actin ratio was decreased by 0.23-fold in C1ORF106 KD cells treated with Y-27632 (*P=*.018) (Fig. 5B), indicating that the increased actin belt thickness in C1ORF106 KD cells is ROCK-dependent. As ROCK can also affect epithelial polarity (21), we inhibited ROCK during spheroid formation, which partially rescued spheroid formation in C1ORF106 KD cells. Indeed, we detected no change in spheroid proportions of controls and observed a 1.53-fold increase in the proportion of spheroids with a lumen in C1ORF106 KD cells (*P=*.0005) (Fig. 5C-D). We also observed decreased proportions of inverted spheroids (0.41-fold; Fig. 5C,E) and of spheroids without a lumen (0.74-fold; *P=*.067; Fig. 5C,F) in C1ORF106 KD cells treated with Y-27632 compared to untreated cells. Strikingly, when we evaluated ROCK1 localization in spheroids without a lumen, we observed that ROCK1 was located in VACs membranes in controls whereas it was excluded from VACs in C1ORF106 KD, further suggesting that the VACs defect in C1ORF106 KD cells is ROCK-dependent (Fig. 5G). Importantly, aPKC, a cell polarity protein (20), is located in VACs, suggesting that the VACs defect in C1ORF106 KD cells could impair cell polarity establishment. As VACs formation is associated with the activation of nmMYO2 around the vesicles, as detected by the phosphorylation of the S19 of the myosin light chain (26), we inhibited nmMYO2 phosphorylation with blebbistatin, which completely rescued VACs accumulation in C1ORF106 KD cells, while we detected no difference in controls (Fig. 6A). Blebbistatin treatment efficacy in all cell lines was confirmed by the impact of the treatment on stress fibers (Fig. S5). These results suggest that the cell polarity defects observed in C1ORF106 KD cells are dependent on an increased activation of ROCK and nmMYO2.

**Figure 5:**
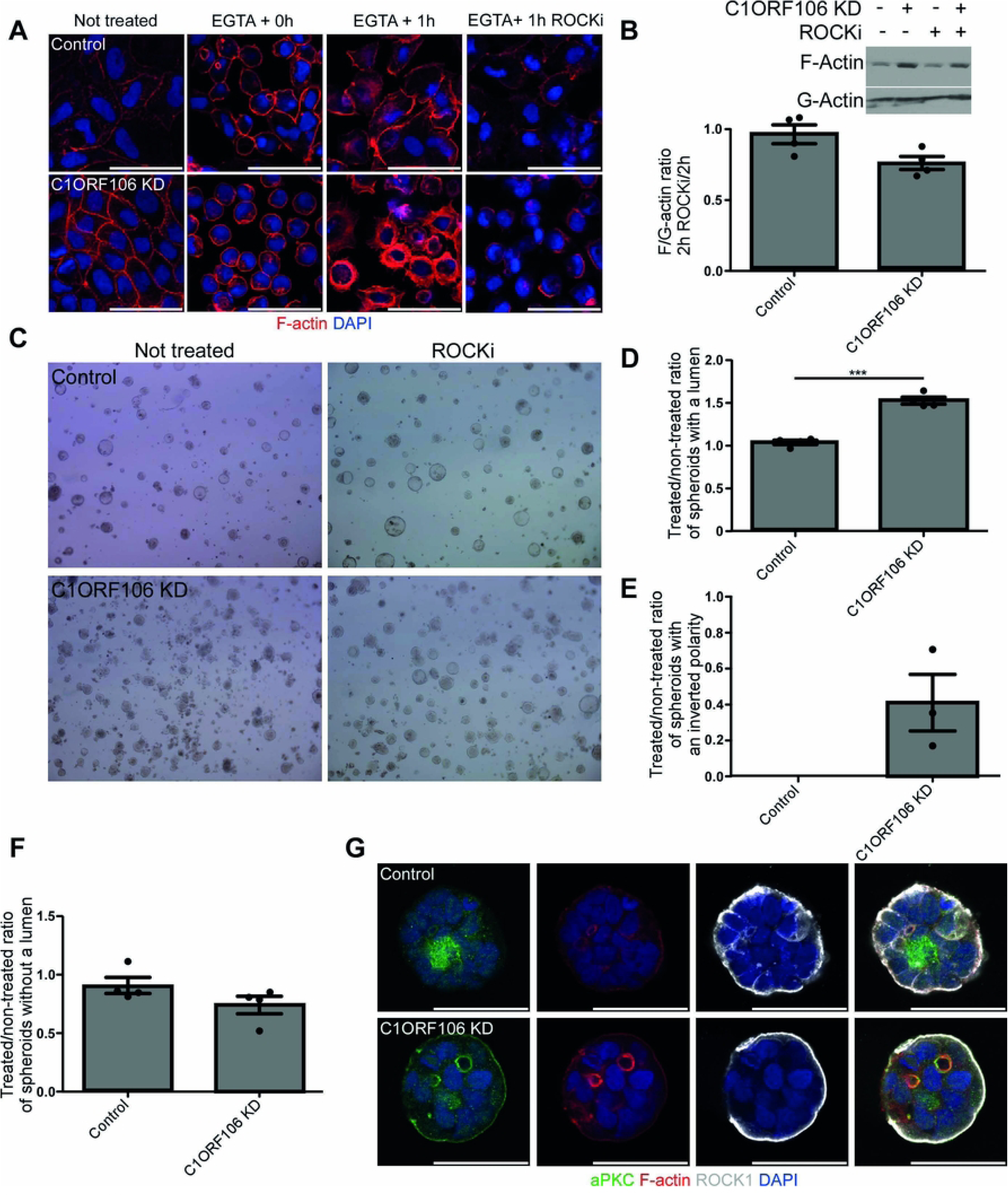
C1ORF106 KD regulates ROCK activation. C1ORF106 KD and control Caco-2 cells were differentiated into a monolayer (A-B) or grown in spheroids (C-G). One sample t-test. Geometric mean + SEM. A-B) Cells were treated with EGTA before being treated with Y-27632 for 1h or 2h. A) Confocal immunofluorescence of F-actin (red) and DAPI (blue). Scale bar=50µm. B) F/G-actin ratio fold change after the Y-27632 treatment. C1ORF106 KD: *P=*.018. n=4. Results are from 4 independent experiments. C-F) Spheroids were grown with or without Y-27632. Brightfield images and quantification of the fold change of each type of spheroids. n=4. D) Spheroids with a lumen. C1ORF106 KD *P=*.0005. E) Spheroids with an inverted polarity. Only the cell lines with at least 1% of spheroids with an inverted polarity were included in the analysis. F) Spheroids without a lumen. \**P=*.067. G) Confocal immunofluorescence of aPKC (green), ROCK1 (grey), F-actin (red) and DAPI (blue) of spheroids without a lumen. Scale bar=50µm.

**Figure 6:**
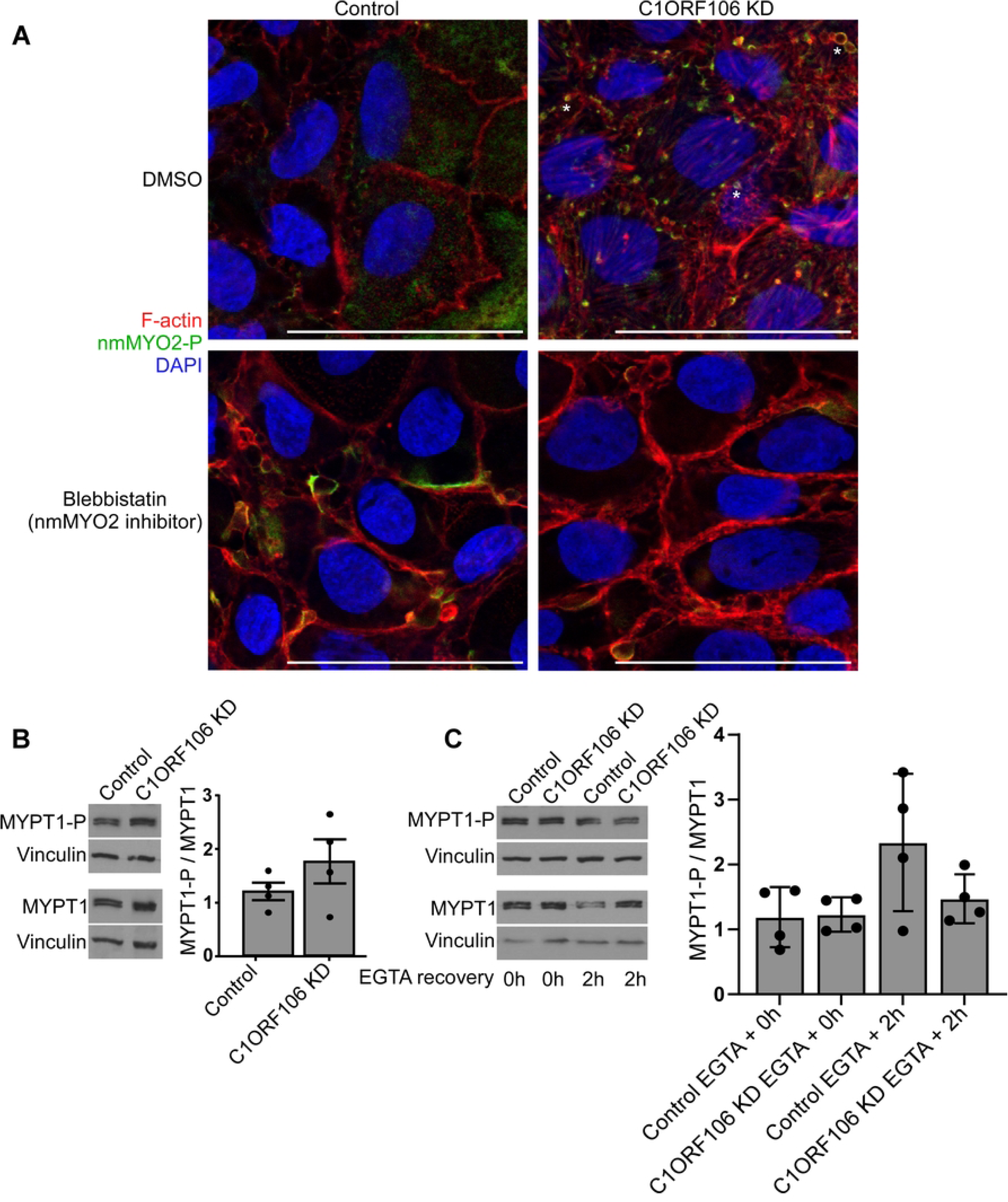
C1ORF106 regulates effectors of ROCK. C1ORF106 KD and control Caco-2 cells were differentiated. A) Cells were treated with nmMYO2 inhibitor blebbistatin or with vehicle (DMSO). Confocal immunofluorescence of F-actin (red), nmMYO2-P (green) and DAPI (blue). Scale bar=50µm. * = Examples of VACs. B-C) MYPT1-P and MYPT1 protein expression was quantified by immunoblot. Geometric mean + SEM. Unpaired t-test with Welch correction. B) n=4. C) Cells were treated with EGTA for 3h and allowed to recover for 2h. n=4.

ROCK also regulates other actin regulators such as MYPT1, which stimulates stress fiber contraction by inhibiting Myosin phosphatase and preventing the dephosphorylation of nmMYO2 (28). We observed that, while not significant, there is a trend of 1.46-fold greater activation of MYPT1 in C1ORF106 KD cells compared to controls (*P=*.075) (Fig. 6B), which is consistent with the increased F/G-actin ratio reported in Fig. S3B and suggests that the alteration in stress fiber dynamics is dependent on the MYPT1/nmMYO2 pathway. However, we detected no significant difference in C1ORF106 KD MYPT1 activation level between 0h and 2h of calcium switch recovery (Fig. 6C), suggesting that the increased actin belt thickness is MYPT1-independent. These results suggest that ROCK is regulating multiple actin-related processes in C1ORF106 KD cells.

### IBD-associated coding variant of C1ORF106 (333F) affects cell polarity

To determine the relevance of these phenotypes to genetic susceptibility in IBD, we studied the impact on cell polarity of the *C1ORF106*-*333^Y/F^* IBD-associated variant (5, 29). We reprogrammed lymphoblastoid cells obtained from two heterozygous patients for the *C1ORF106*-*333*^Y/F^ variant and from two individuals carrying the wildtype allele (333^Y/Y^) to generate hiPSC lines. We validated the genotype status of these hiPSC lines for the the *C1ORF106*-*333*^Y/F^ variant via Sanger sequencing and characterized these hiPSC lines for chromosomal rearrangements, their capacity to differentiate into the three germ layer and their expression of pluripotency markers (Fig. S6) We then differentiated them into intestinal epithelial lines, growing them as intestinal epithelial spheroids and characterizing these spheroids (Fig. S7). Similarly to C1ORF106 KD, the proportion of spheroids with a lumen was lower (33%) in *C1ORF106*-*333*^Y/F^ cell lines than in controls (*C1ORF106*-*333*^Y/Y^) (87%) (Fig. 7A, S8) (*P=*.0243), suggesting that this variant affects spheroid formation. Ezrin was located on the apical side of the spheroids with a lumen and was located in vesicles in spheroids without a lumen (Fig. 7B). Interestingly, nmMYO2-P was mainly located on the apical side of the spheroids with a lumen from *C1ORF106*-333^Y/Y^, while it seemed located on both the apical and basal sides in *C1ORF106*-333^Y/F^ spheroids (Fig. 7C, S9), suggesting that nmMYO2 is dysregulated in these cells.

**Figure 7:**
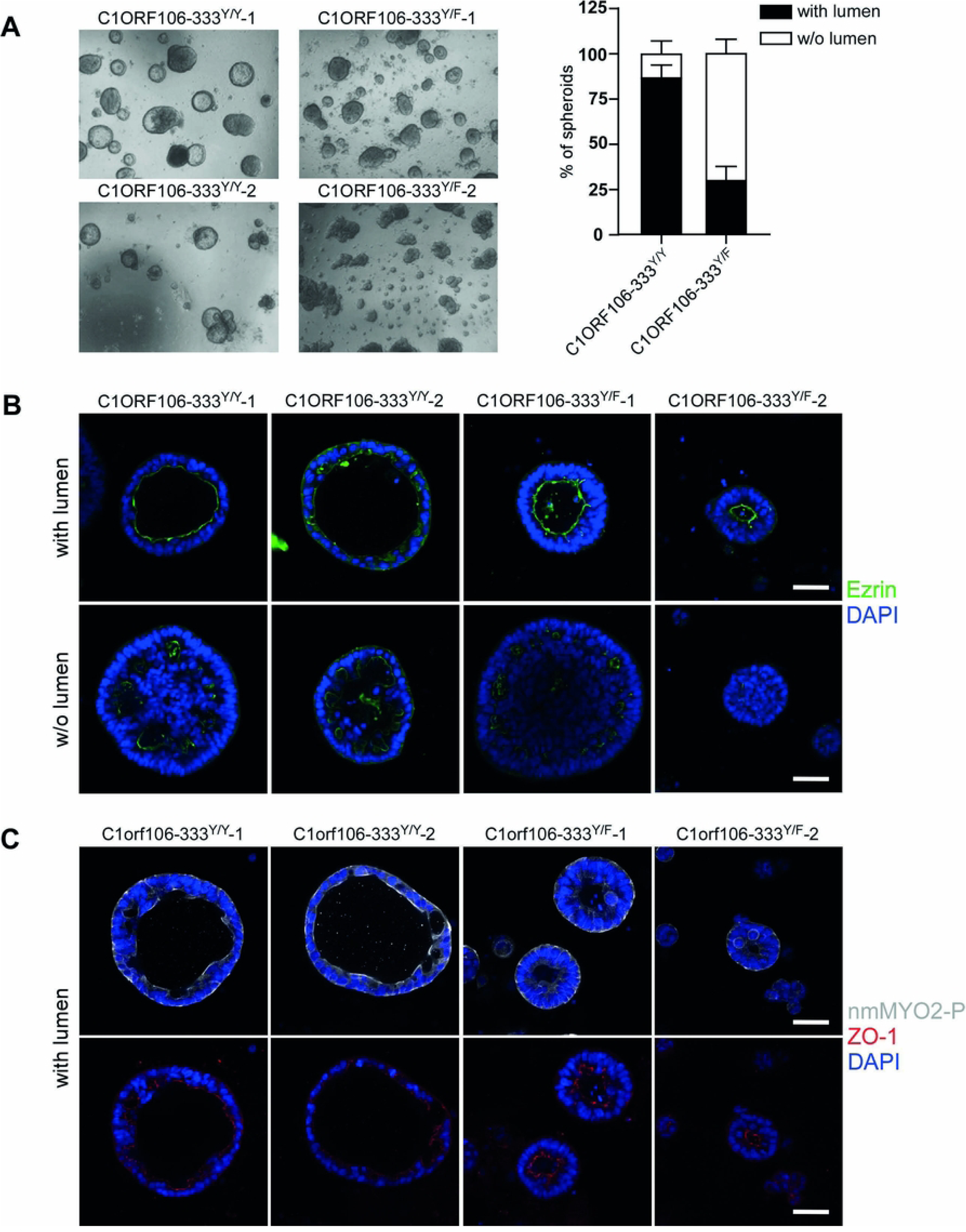
C1orf106*333F variant regulates cell polarity. hiPSC-derived Intestinal epithelial spheroids were grown for 10 days. A) Brightfield images and quantification of each type of spheroids. n=2. Geometric mean + SEM. Welch two sample t-test. *P=*.024. Results are representative of 2 independent experiments. B-C) Confocal immunofluorescence microscopy of DAPI (blue), B) Ezrin (green) and C) ZO-1 (red) and nmMYO2-P (white). Scale bar=50µm.

## Discussion

Until now, IBD therapeutics have focused on regulating immune cell functions rather than targeting defects in epithelial permeability (30). Studies elucidating epithelial functions important in IBD are emerging and are highlighting an opportunity to develop new treatments with novel mechanisms of action (30, 31). Importantly, the knowledge of a genetic mechanism of disease development increases the likelihood that a drug targeting this pathway will be efficient in treating this disease, supporting the importance of studying new mechanisms of action of IBD-associated genes (32). C1ORF106 has previously been reported to control epithelial permeability through the regulation of TJ and AJ (5, 6). We have focused our efforts to understand this pathway because increased intestinal epithelial permeability in asymptomatic individuals is associated with an increased risk of developing CD (3). We used C1ORF106 KD Caco-2 cells to demonstrate the impact of a decreased expression of C1ORF106 on TJ formation, cell polarity establishment and actin contraction. This model is relevant because it was previously demonstrated that the IBD-associated *C1ORF106* coding variant Y333F decrease protein stability by ∼40% (5).

Herein, we have demonstrated that C1ORF106 regulates the actin dynamics during apical junction formation and that the epithelial barrier recovery is altered in IEC with reduced C1ORF106 expression. C1ORF106 was previously shown to regulate the AJC through the regulation of AJ stability and ZO-1 localization, an adaptor protein linking the TJ to the actin belt (5, 6, 33). This interaction is important for the regulation of epithelial permeability by the actin belt contraction as demonstrated by the impact of TNF-α on actin belt contraction and barrier integrity (15). Kinases such as ROCK and MLCK, which can be activated by pro-inflammatory cytokines and regulate actin belt contraction are implicated in the regulation of epithelial permeability (15, 33). Notably, similarly to C1ORF106 KD cells, IFN-γ regulates the internalisation of AJC proteins into VACs by regulating the actin belt contractility through ROCK (34). As pro-inflammatory cytokines are increased in IBD patients(30), these results suggest that the decrease in C1ORF106 expression in susceptible individuals could increase ROCK activity and make them more sensitive to pro-inflammatory cytokines.

We also demonstrated that other ROCK-dependent processes like stress fiber regulation are altered in C1ORF106 KD cells. We observed an impaired F-actin regulation without reorganization of focal adhesions as expected with a dysregulation of nmMYO2 (18, 35). Importantly, an impaired cell adhesion can lead to epithelial cell detachment and inflammation as observed in mice depleted for FERMT1, an IBD-associated gene regulating cell adhesion, where IEC detachment induced an important cytokine production and immune cell recruitment (36). Taken together, this suggest that a disturbed stress fiber regulation in C1ORF106 KD cells could facilitate infections and cell detachment, which would stimulate inflammation.

Importantly, an increased actin contraction can impair cell polarity establishment, which controls the localization of receptors and transporters regulating barrier permeability (21, 24). We observed an altered cell polarity establishment characterized by a decreased ratio of spheroids with a lumen and by an altered localization of apical markers in C1ORF106 KD spheroids and in patient-derived *C1ORF106*-*333*^Y/F^ spheroids generated from hiPSC cells. We demonstrated that ROCK inhibition during spheroid formation partially rescued the impaired spheroid formation of C1ORF106 KD cells, indicating that this process is dependent on actin contraction. An increase in ROCK activation can impair the asymmetric matrix protein secretion, which impairs the signaling pathways regulating cell polarity establishment (21). This is further supported by recent work demonstrating that HCT8 spheroids overexpressing *C1ORF106* and treated with an inhibitor of ROCK had an increased barrier function through a CYTH2-dependent mechanism (9). Taken together, these results demonstrate that the cell polarity impairment in C1ORF106 KD cells is linked to ROCK-dependent F-actin regulation. Moreover, our results suggest that the alteration in spheroid formation is associated with the accumulation of VACs, which is dependent on nmMYO2 activation (24), and which happens before RAB11 recruitment to these vesicles. Interestingly, defects in VACs formation are similar to microvillus inclusions that have been observed in Microvillus Inclusion Disease (MVID), an enteropathy associated with an internalization of brush border components resulting in severe diarrhea (25). MVID is characterized by a loss of IEC polarity leading to an altered localization of transporters involved in nutrient absorption (25). Defects in the interaction between RAB11 and MYO5B have been associated with the formation of microvillus inclusions in mouse models (37). These results strongly suggest that VACs have an important role in intestinal barrier function by regulating cell polarity.

We demonstrated in C1ORF106 KD cells that cell migration, a process important for wound healing (30), is faster, consistent with a previous study in C1ORF106 KO mice organoid-derived epithelial cells (5), and that it is less directional. Importantly, a loss of cell polarity can impair the capacity of cells to migrate persistently in the direction of the wound (38). Leader cells at the migration front dictate the migration orientation and cells migrating independently migrate faster and in a less coordinated manner (39). Interestingly, we observed a disorganization of the migration front structure. AJ regulate the integrity of groups of migrating cells and the coordination of the migration of adjacent cells by inducing a contact inhibition of locomotion in the directions in which there are cells (39). When the junctions are intact, the leader cells migrate in the only free direction (39). As C1ORF106 regulates apical junction formation and stability, this could explain the impaired migration orientation and coordination that we observed in C1ORF106 KD cells.

Overall, we demonstrated that C1ORF106 regulates AJC through its impact on F-actin dynamics in IEC. This process is dependent on ROCK activation and impacts on multiple F-actin functions like cell constriction and cell polarity. An impaired AJC formation and stability could increase the passage of molecules between the cells and the downstream recruitment and activation of the immune system. We also demonstrated an impact of the IBD risk allele *C1ORF106*-*333*^Y/F^ on cell polarity, demonstrating the importance of this pathway in IBD. Interestingly, multiple ROCK inhibitors have been approved in humans to treat other diseases like cerebral vasospasm and glaucoma (28, 40). This pathway represents an interesting new target for drug development in IBD focused on reestablishing epithelial barrier function although more studies would be needed as studies reported conflicting results on the impact of ROCK inhibition on epithelial permeability (19, 41). As these inhibitors have a systemic impact including changes in heart rate and in lymphocyte counts (40), it would be crucial to develop a treatment that acts locally on the gut. Globally, our study demonstrates that C1ORF106 regulates not only the AJC, but also epithelial cell structure and migration, which makes it an important regulator of intestinal barrier function.

## Abbreviations

AJ: Adherens junctions
AJC: Apical junctional complex
CD: Crohn’s disease
FBS: fetal bovine serum
hiPSC: human induced pluripotent stem cell
IBD: Inflammatory bowel disease
IEC: Intestinal epithelial cells
KD: Knock-down
KO: Knock-out
MVID: Microvillus Inclusion Disease
nmMYO2: non-muscle myosin-2
Penicillin and streptomycin: P/S
ROCK: Rho Associated Coiled-Coil Containing Protein Kinase
RT: Room temperature
SEM: standard error
TEER: transepithelial electrical resistance
TJ: Tight junctions
UC: Ulcerative colitis
VACs: Vacuolar apical compartments

## Acknowledgements

IHM received scholarships from the Claude-Lise Richer funds of the Université de Montréal, from the Biomedical Science program of the Université de Montréal, from the Famile Valiquette Research funds of the Université de Montréal and from the Anciens de Shawinigan foundation. This work was funded by the Canadian Institutes of Health Research (Genetics; Nutrition, Metabolism and Diabetes; Infection and Immunity; CIHR#451128), by the National Institutes of Diabetes, Digestive and Kidney Diseases (UO1DK062432) and by the IMAGINE-SPOR chronic disease network Canadian Institutes of Health Research (Infection and Immunity; CIHR#145105).

## Data availability statement

All relevant data are within the manuscript and its Supporting Information files.

## Notes

### Competing Interest Statement

The authors have declared no competing interest.

